# The global communication architecture of the human brain transcends the subcortical - cortical - cerebellar subdivisions

**DOI:** 10.1101/2023.07.07.548139

**Authors:** Julian Schulte, Mario Senden, Gustavo Deco, Xenia Kobeleva, Gorka Zamora-López

## Abstract

The white matter is made of anatomical fibres that constitute the highway of long-range connections between different parts of the brain. This network is referred to as the brain’s structural connectivity and lays the foundation of network interaction between brain areas. When analysing the architectural principles of this global network most studies have mainly focused on cortico-cortical and partly on cortico-subcortical connections. Here we show, for the first time, how the integrated cortical, subcortical, and cerebellar brain areas shape the structural architecture of the whole brain. We find that dense clusters vertically transverse cortical, subcortical, and cerebellar brain areas, which are themselves centralised by a global rich-club consisting similarly of cortical and subcortical brain areas. Notably, the most prominent hubs can be found in subcortical brain regions, and their targeted in-silico lesions proved to be most harmful for global signal propagation. Individually, the cortical, subcortical, and cerebellar sub-networks manifest distinct network features despite some similarities, which underline their unique structural fingerprints. Our results, exposing the heterogeneity of internal organisation across cortex, subcortex, and cerebellum, and the crucial role of the subcortex for the integration of the global anatomical pathways, highlight the need to overcome the prevalent cortex-centric focus towards a global consideration of the structural connectivity.

## INTRODUCTION

Since the first maps of brain-wide white matter communication pathways in the mammalian brain for the cat and the macaque brains (Scannell et al., 1995; Scannell & Young, 1993; Young, 1993), in the last three decades, the field of brain connectivity has emerged to finally shed light about the puzzling function of the white matter. In these years we have learned that the white matter forms a complex network of communication that determines how different brain regions interact with each other. While 20th-century neuroscience described the localization and the specialisation of brain regions for the processing of specialised tasks, brain connectivity has opened up a picture of the brain in which those specialised regions are very much interdependent. This interconnectivity is not only fundamental to understanding brain functioning in a healthy state but also in the diseased brain as alterations to this network are related to various psychiatric and neurological diseases (Kim et al., 2014; Qiu et al., 2010; Shao et al., 2012; Skudlarski et al., 2010).

Due to this importance, many studies have investigated the cortical structural network discovering general architectural properties. Among other findings, the cortex has been found to be segregated into communities i.e. clusters of brain areas that are more densely connected with each other than outside of that community (Betzel et al., 2013; Hilgetag et al., 2000; Hilgetag & Kaiser, 2004; Newman, 2004; Zamora-López et al., 2011). These communities are structurally integrated by widely connected brain regions (i.e. hubs), which themselves form a densely interconnected supra-structure referred to as rich-club (Senden et al., 2014, 2017, 2018; van den Heuvel & Sporns, 2011; Zamora-López, 2010, 2009; Zhou & Mondragon, 2004). Intriguingly, cortical rich-club nodes can be found in all resting-state functional networks and therefore these highly interconnected hubs might serve as an anatomical substrate for structural-functional integration (van den Heuvel & Sporns, 2013).

Despite this success, the studies of brain connectivity have helped overall to promote a cortex-centric view of the human brain. The integration of the subcortical and cerebellar regions into network analyses has been rather anecdotal. Typically, the studies of human brain connectivity have included a few subcortical regions but at a low level of resolution, e.g. considering the (left/ right) thalamus or hippocampus as a single region. This research, as observed for the cortex alone, also demonstrated small-world properties showing that despite a large network, brain regions are closely connected (i.e. short average path length) and tend to cluster together (Gong et al., 2009; Hagmann et al., 2008; Iturria-Medina et al., 2008; Sporns & Zwi, 2004). Moreover, several subcortical brain areas were found to be part of the rich-club (van den Heuvel & Sporns, 2011). Less research has been dedicated to the cerebellum individually, where a modular network architecture with small-world characteristics has also been found (Kim et al., 2014).

However, the cortex, the subcortex, and the cerebellum are tightly interlinked via various pathways, which are crucial for both cognitive and sensory functions. Hence, it is fundamental that we understand how they synergistically shape the global architecture and how structural integration is achieved. To answer this question, we present here the first network analysis of the human connectome that comprises the cortex, the subcortex, and the cerebellum altogether.

We find that the global network displays a modular and hierarchical organisation – similar to the one typically described for the cortex alone. However, a subsequent community detection reveals a modular organisation that transcends the classical – cortical, subcortical, cerebellar – subdivision pointing to functional communities that encompass regions of all three components. Notably, these communities are integrated by a global rich-club, which at its core is subcortically dominated. Congruently, subcortical hubs represent the most connected brain regions in the global network, and targeted in-silico lesions to them proved to be most harmful for global signal propagation. These results highlight the centralising role of subcortical hubs and suggest a crucial role for global brain functioning. Moreover, similarly to the cortex alone (van den Heuvel & Sporns, 2013), we also find nearly all functional resting-state networks being represented in the global rich-club, indicating that it not only serves structural but also functional centralization. Finally, we can observe both similarities and differences in the network architectures of the global, cortical, subcortical, and cerebellar (sub)networks. While all networks exhibit a near-optimal short-average path length and information transmission capacity, they display distinct variations in degree distribution, clustering, and the propensity for small and high-degree areas to connect with one another. The heterogeneity among these (sub)networks and the substantial role of the subcortex in global structural centralization emphasise the importance of transitioning from a cortex-centric perspective to a comprehensive analysis of the entire structural network. This study is also bound by the imaging resolution and the applied parcellation. More detailed circuitries, which are certainly also important to understand, will be coming in the near future thanks to the upcoming high-resolution connectomes. This paper represents a first step delineating the general architecture of the brain in its holistic perspective.

## RESULTS

The goal of this study is to uncover the brain-wide organization of the communication network established by the long-range white matter connections in the human brain, encompassing both the cerebral cortex, the subcortical regions, and the cerebellum. Therefore, we study the group-averaged structural connectivity (SC) matrix extracted via diffusion-weighted imaging for 32 healthy human volunteers of the Human Connectome Project. We parcellated the brain-wide network following Ji et al. (2019), which divides the brain into n = 718 regions of interest (ROIs), of which n_cor_ = 360 correspond to cortical regions, n_sub_ = 233 subcortical and n_cer_ = 125 cerebellar ROIs. We simultaneously investigate the global network and its individual cortical, subcortical, left, and right cerebellar subnetworks. Given empirical evidence that all connections from the cerebellum to the cortex are mediated via contralateral connections through the Pons, and the Thalamus (D’Angelo, 2018; Palesi et al., 2015, 2017; Ramnani, 2006), we filtered out direct cortico-cerebellar connections and also ipsilateral connections from the cerebellum to the thalamus and brainstem. Similarly, direct cerebellar inter-hemispheric connections were also filtered as the cerebellar hemispheres are believed to be indirectly connected via the vermis (D’Angelo, 2018) and we therefore separately investigated the individual hemispheres of the cerebellum.

The final global network included the cortical regions according to Glasser et al. (2016) and a subcortical parcellation according to Ji et al. (2019) where all parcels encompassed major subcortical/cerebellar structures as defined by Freesurfer. Subcortical ROIs included bilaterally the nucleus accumbens (n=13), brainstem (n=47), caudate nuclei (n=17), diencephalon (n=40), hippocampus (n= 29), globus pallidus (n=20), putamen (n=18), thalamus (n=38), amygdala (n= 11), and 125 left and right ROIs encompassing the cerebellar cortex (see Methods).

We firstly performed a system-level analysis investigating how the cortex, subcortex, and cerebellum are globally interlinked with each other and whether non-apparent internal organisation could be found in the form of network communities. Next, we explored the structural centralization of these communities. We identified the primary locations of brain hubs and examined rich-club properties. Subsequently, we examined the (sub)networks’ vulnerability towards targeted and random brain lesions and lastly performed a more detailed network characterization of the global and individual (sub)networks.

### Brain-wide communication architecture transcends the anatomical cortical, subcortical, and cerebellar division

We begin our investigation by revealing the systems-level picture of the brain-wide network. Therefore, we first study the density of connections within each of the brain’s subdivision – cortex, subcortex, cerebellum – and the amount of cross-connections between them. Second, we apply a community detection method to reveal the modular division of the brain-wide network based solely on the organization of the underlying white-matter communication pathways. Finally, we compare the similarities and differences between the divisions.

The density of connections in a (sub)network indicates the fraction between the number of links present in the (sub)network and the maximum number of connections possible. Considering the anatomical division of the brain, internal densities ranged from 23.4% for the cortex to the denser 32.4% for the subcortex, Figure 1A. We also found comparable large cross-connection densities – approximately 24% – between the cortex and subcortex, and 19.3% between the subcortex and the cerebellum. The cortico-cerebellar connection and cerebellar interhemispheric connections were a priori filtered out, as these are not mirrored by anatomical literature (D’Angelo, 2018; Palesi et al., 2015, 2017; Ramnani, 2006). These observations underline that a heavy intercommunication exists between the cortex, the subcortex, and the cerebellum. Moreover, these anatomical subdivisions do not seem to form clearly separated communities from a pure network perspective as the cortical and the cerebellar regions have approximately the same probability to connect to themselves as to subcortical regions.

**Fig. 1:**
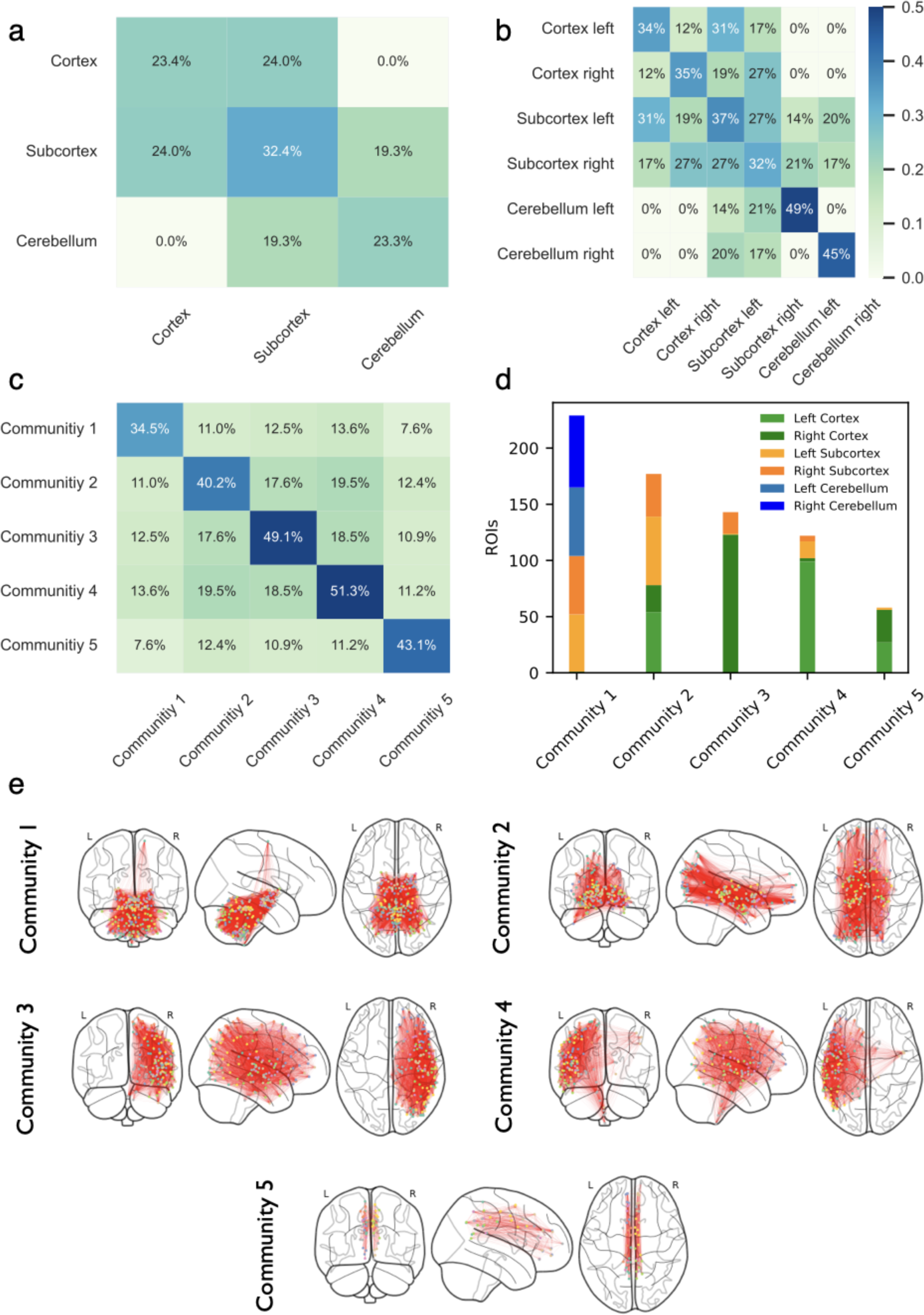
System-level connectivity between brain areas and modules. **a,b** Connection densities within and between the (left & right) cortex, subcortex, and cerebellum. High interconnectivity between the cortex, subcortex, and cerebellum can be observed. **c** Connection densities within and between the detected communities. These showed low densities between each other and equally or even higher internal densities than the cortex, subcortex, and cerebellum. **d,e** Regional and spatial distribution of the communities. Communities 1-4 vertically stretched over cortical, subcortical, and cerebellar boundaries. Both lateralized (communities 3&4) and more medially arranged communities (1,2, and 5) were detected.

Splitting up the brain-wide network into left and right hemispheres, different lateralisation patterns were detected, Figure 1B. Regarding the interconnections within each of the anatomical subdivisions, the cortex, and the subcortex display distinctive intra-/inter-hemispheric connection densities. For cortical regions, it is about three times more likely to connect to other cortical regions in the same hemisphere (∼35%) than in the contralateral hemisphere (12%). Similarly, ipsilateral cortico-subcortical connections are more likely than contralateral connections (∼29% ipsilateral vs. ∼18% contralateral). Whereas subcortical brain regions showed a less pronounced lateralization, the inter-hemispheric connections of the cerebellum, and cortico-cerebellar connections were not present as these were a-priori filtered out (D’Angelo, 2018; Palesi et al., 2015, 2017; Ramnani, 2006).

As the cortex, subcortex, and cerebellum are tightly interconnected and no distinct differentiation was observed, could the global connectome hide another non-apparent internal organisation? To investigate this possibility, we applied a community detection algorithm to find structural communities. Generally, a community is understood as a grouping of nodes, which are more densely connected inside than outside of that community (Newman, 2004). For that, we used the Leidenalg algorithm (Traag et al., 2019) together with the Reichardt & Bornholdt (2006) modularity function to evaluate the goodness of the partition. This approach integrates a resolution parameter γ that influences the total number and the size of the communities encountered. The algorithm was run in the range of γ ∊[0.6..1.9] with 100 realisations per γ. The optimal solution was identified for γ = 1 with a modularity value of Q = 0.232, dividing the network into a partition of five communities. Notably, these were characterised by high internal densities (34.5% - 51.3%) comparable to – or even higher than – the internal densities of the cortex (23.4%), subcortex (32.4%), and cerebellum (23.3%; Fig. 1c,a). Also low inter-community densities can be observed highlighting the distinctiveness of the communities. This implies that the communities are better segregated than the anatomical subdivision.

Notably, all five communities were characterised by different regional and hemispherical structural profiles demonstrating that dense groupings of brain areas can be found across cortical, subcortical, and cerebellar boundaries (Fig. 1d,e). The first community evenly comprised left and right ventrally located subcortical and cerebellar ROIs. Similarly, the second community also comprised ROIs from both hemispheres but encompassed cortical and subcortical ROIs. The third community consisted primarily of right hemispherical ROIs in the cortex and subcortex. The fourth community is the counterpart to the third as it mainly concerned left cortical and subcortical ROIs. Interestingly, the ROIs in communities three and four were more laterally located, had also contralateral cortical ROIs, and were also connected to deeper subcortical ROIs. Community five, which is the smallest one, is more medially located and evenly comprised of left and right cortical ROIs. Whereas this community mainly encompassed cortical ROIs, some subcortical ROIs could be found here as well. Although their functionality is not clear at this point, communities three and four could support more lateralized cognitive processes such as language (Knecht et al., 2000; Olulade et al., 2020) and attention (Bartolomeo & Seidel Malkinson, 2019) as for instance shorter average path lengths, as well as different information processing. Congruently, communities one and two might support more distributed neural processes and support information integration due to their central arrangement. Community five could also support these integrative processes, however, mainly focused on cortical integration. Interestingly, communities one to four, encompassing most ROIs in the network, vertically transverse cortical, subcortical, and cerebellar boundaries, and consequently offer a differential perspective on global structural connectivity as compared to the traditional anatomical division.

### The core of the global rich-club is subcortically dominated

The community detection revealed global communities spanning over cortical, subcortical, and cerebella ROIs. However, it is not clear how these system-wide communities might be structurally integrated. Hub regions usually play a crucial role for structural centralisation. We identified the most connected brain regions in the global network. Subcortical hubs, particularly ROIs in the brainstem, putamen, diencephalon, and globus pallidus are connected to more than 500 other brain regions and have consequently a crucial centralising role (Fig. 2a). The 10 most connected brain regions could reach around ∼94% of the global network (Fig. 2b). Expanding the analysis and considering the 72 most connected brain regions (10% of the total) confirms the clear dominance of subcortical brain regions (i.e. 68%; Fig. 2e), which could reach around ∼97.9% of the total network (Fig. 2b). Further investigating how these hubs split between the detected communities, we observed that a larger fraction of these hubs is found in the interhemispheric communities (1 and 2; Fig. 2e). This was congruent with the structural profile of those communities as they comprised to a large extent subcortical brain regions.

**Fig. 2:**
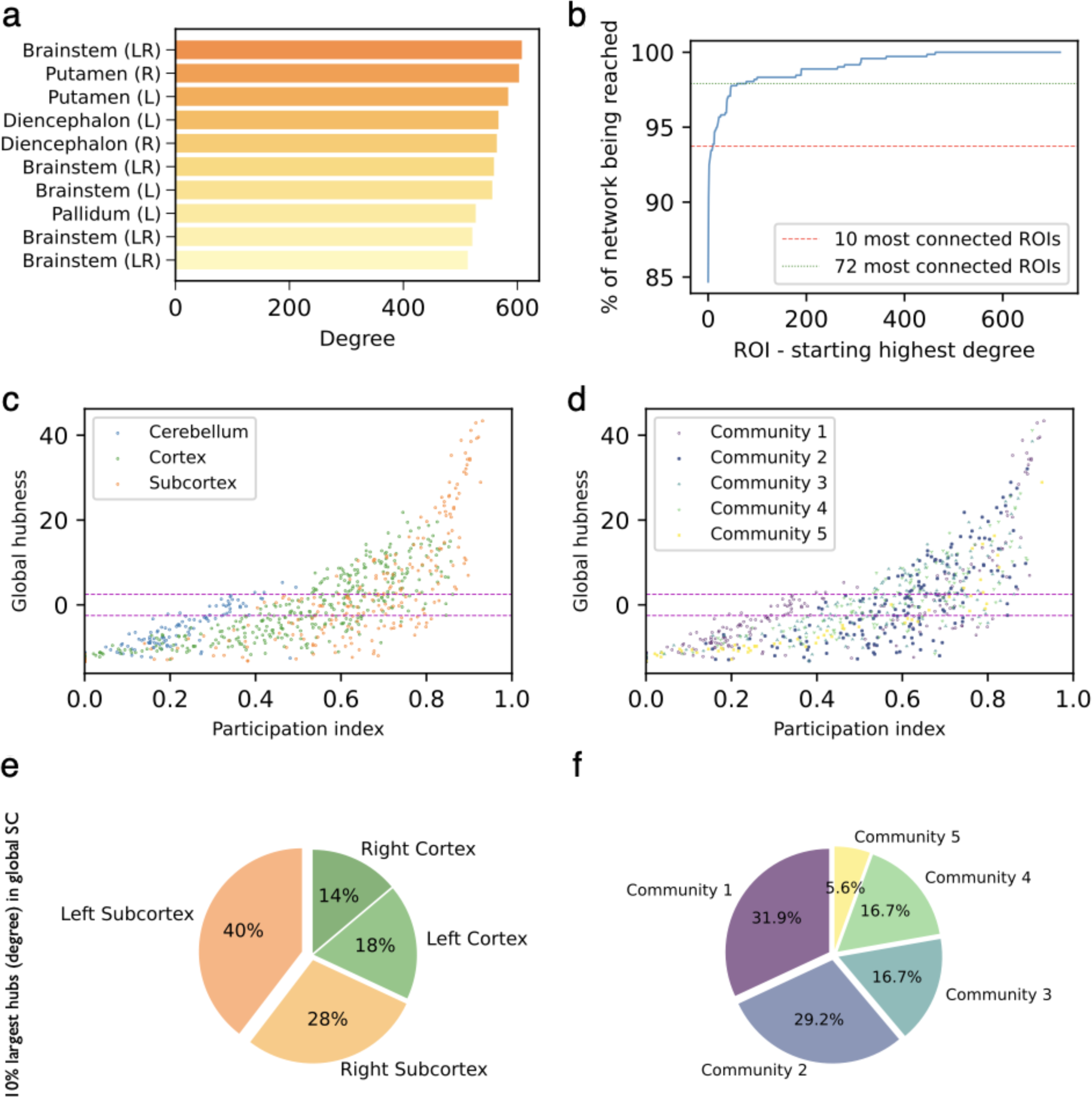
Leading integrative hubs were found in the subcortex. **a** 10 most connected ROIs in the global network (n=718) **b** Global network reach iteratively combining nodes starting with the most connected hub. Already the 10 most connected hubs can reach 93.7% and the 72 most connected nodes 97.9% of the entire network. **c,d** Global hubness and participation coefficient of all nodes in the global network. Leading hubs were found in the subcortex (and in communities 1-5) and showed a high participation coefficient underlining their integrative properties. The horizontal red lines indicate the range of a random graph, where 99% of nodes are located (i.e. global hubness larger than (-)2.5) **e,f** 10% of largest hubs (72 ROIs) in the global network measured by their degree. Around 68% of them are found in the subcortex and 61% of them in communities 1 and 2.

To provide a broader overview of how well brain regions are generally connected, we looked at the global hubness *h_i_*, which compares the degree *k_i_* of a brain region *i* with the degree distribution of a corresponding random network of equal size *N* and density *ρ*, thus creating a common statistical benchmark (Klimm et al., 2014). Hub regions (i.e. *h_i_* > 2.5; Fig. 2c,d) were mainly found in the cortex, and subcortex, and some also in the cerebellum. In contrast, all five communities showed hub regions (Fig. 2d,f). To further evaluate how hubs link to the identified communities, we looked at the participation coefficient *ρ_i_*, which measures how uniformly a node *i* distributes its links across the communities. A participation coefficient of zero indicates that a node has only links within its own community, whereas a value of one indicates a uniform distribution of links towards all communities (Klimm et al., 2014). Generally, for the most prominent hubs a high participation coefficient was observed showing that these brain regions structurally centralise the global communities. In detail, these hubs uniformly distributed their links to the communities and consequently acted as overarching integrators rather than being distinguishable members of a single community. Especially subcortical but also cortical hubs had this structural property underlining their importance for global information exchange. In contrast, the few cerebellar hubs showed a low participation coefficient indicating a local reach. Cross-community integration of cerebellar information is therefore indirectly achieved by subcortical hubs in community one acting as centralising bridges. This is also replicated by anatomical evidence as the cerebellum is connected to cortical brain regions via the thalamus and pons (D’Angelo, 2018; Palesi et al., 2015, 2017; Ramnani, 2006).

The observation that subcortical and cortical hub regions have a global reach and cannot be clearly attributed to individual communities pointed toward a structural principle previously observed in the cortex: it has been shown that structural communities are centralised by hubs, which are themselves densely interconnected with each other (Zamora-López, 2010, 2009). This densely connected supra-structure is referred to as rich-club (Zhou & Mondragon, 2004), and while communities can be found spanning over the cortex, subcortex, and cerebellum it is not clear whether they are also centralised globally by a rich-club. To determine whether hubs are highly interlinked with each other, we calculated the k-density across different values of k. The k-density measures the internal density of connections between the hubs, such that k′ > k. At the same time, the observed rich-club properties were compared to a random graph ensemble for the individual networks (1000 surrogate networks for each network e.g. global, cortical, etc.) to evaluate how much of the observed characteristics could be explained purely by the networks′ degree distributions. The cerebellar structural network was separately investigated for the left and right hemispheres as both hemispheres are believed to be indirectly connected via the vermis. Rich-club properties could be observed for all (sub)networks as their hubs (high degree nodes) were tightly interlinked (i.e. high k-densities; Fig. 3a). Whereas the global and cortical structural (sub)networks showed a higher k-density between hubs than expected from their degree distributions, the subcortical, left, and right cerebellar subnetworks showed no significant deviation.

**Fig. 3:**
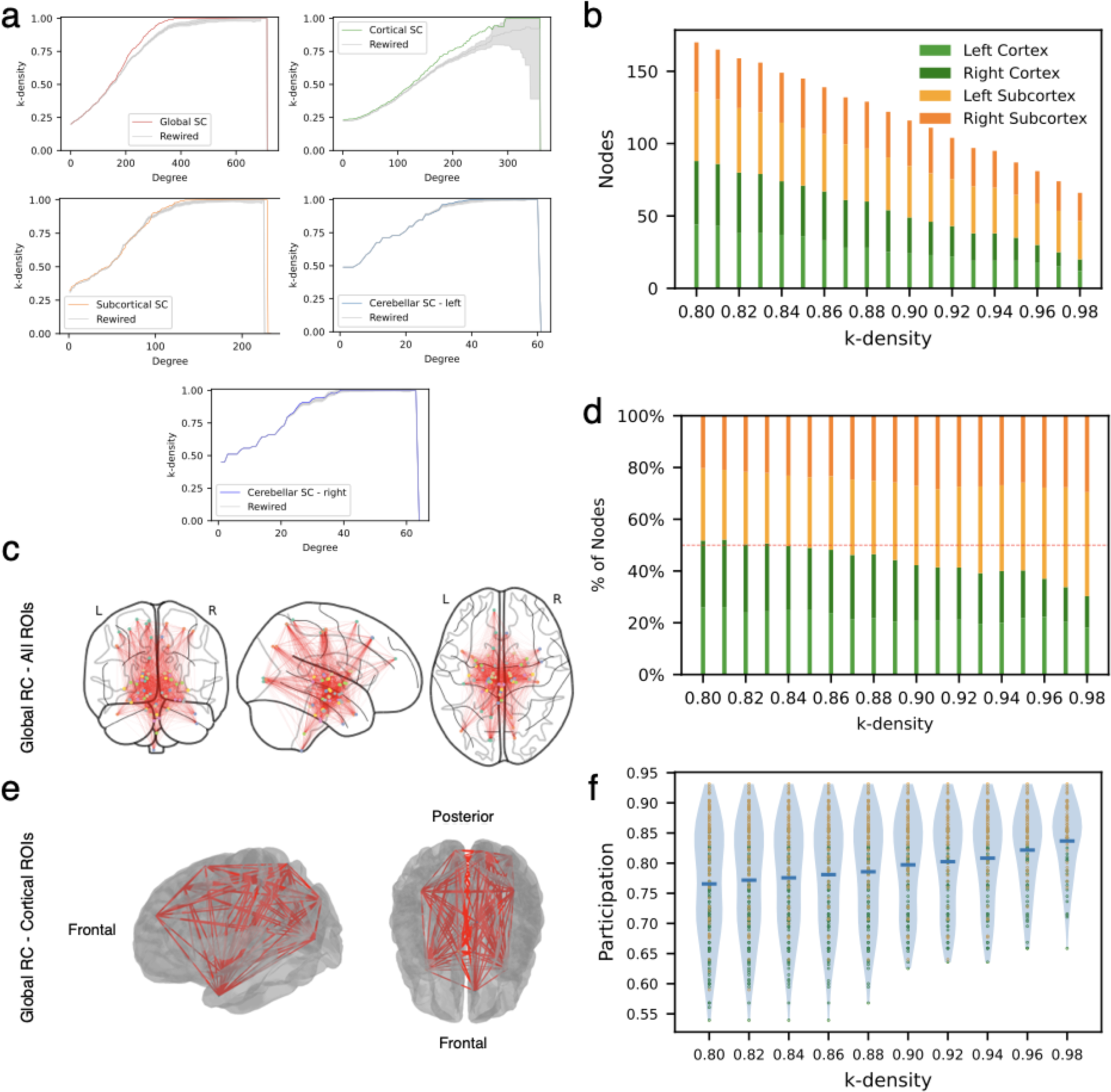
Rich-club properties of the global and individual (sub)networks. **a** k-density charts for all (sub)networks. The silver line represents the mean of a random graph ensemble (n=1000) with the shaded lines 2 standard deviations. All (sub)networks showed rich-club properties as their hubs (high-degree nodes) were tightly interlinked. For the global and cortical (sub)networks these high densities could not be explained by the networks’ degree distribution. **b,d** Total number (b) and percentages (d) of nodes in the global rich-club per k-density. The global rich-club consisted of cortical and subcortical ROIs. At lower k-densities cortical and subcortical ROIs equally participated in the rich-club, whereas at the core (i.e. large k-densities) the subcortical ROIs represented the majority. **c** Spatial allocation of all global rich-club nodes at a density of 98% (66 ROIs). The rich-club is mainly medially located along the longitudinal fissure. **e** Only cortical ROIs of the global rich-club. Nodes can be observed located at all brain lobes. **f** The participation coefficient of all global rich-club nodes per k-density. Generally, subcortical ROIs (orange) had a larger participation coefficient than cortical ROIs (green).

For the global rich-club, it can be observed that it was characterised evenly by subcortical and cortical hubs and no cerebellar brain regions (Fig. 3b,d). Notably, subcortical brain regions represented a majority at the core (i.e. high internal densities) of the rich-club. For instance, at a k-density of 0.98, 70% of the rich-club consisted of subcortical ROIs (n=46) encompassing generally regions in the brainstem (n=12), diencephalon (n=8), thalamus (n=10), amygdala (n=5), hippocampus (n=3), globus pallidus (n=2), putamen (n=4), and caudate nucleus (n=2). Cortical rich-club ROIs (n=20; 30%) stretched over the whole cortical surface encompassing all cortical lobes (Fig. 3c). Interestingly, most cortical and subcortical rich-club ROIs were medially located proximal to the longitudinal fissure. Figure 3f shows that rich-club regions, especially subcortical ROIs (orange), were characterised by a high participation coefficient and thus centralised the different communities. These findings underline the importance to consider the entire connectome as a global rich-club can be found, which centralises the detected communities with subcortical hubs serving as especially integrative hubs.

### Global structural rich-club centralises functional networks as well

The global rich-club forms the basis for structural centralization. However, this raises the question of how this architectural property relates to global functional network properties. Previous work showed that the presence of a rich-club architecture increases a network′s diversity of functional repertoire and that it may achieve functional integration by exhibiting oscillations and harmonising asynchronous brain regions (Senden et al., 2014, 2017). Congruently, it is suggested that the cortical rich-club might serve as a structural backbone for cross-linking functional resting-state networks as rich-club nodes were present in all of these networks (van den Heuvel & Sporns, 2013). To investigate this interplay, we analysed how the global structural rich-club relates to cortical-subcortical functional network organisation. Here, the functional networks are based on the community detection of Ji et al. (2019) performed on fMRI data (see Methods).

The rich-club representation measures the percentage of nodes in a functional network being represented in the global rich-club. Independent of the internal rich-club densities, nearly all functional networks except the ventral multi-modal network participated in the global rich-club (Fig. 4a) mirroring the previous cortical findings by van den Heuvel and Sporns (2013). This could be an indication that the global rich-club facilitates structural-functional integration. Interestingly, no clear majority of any network can be observed at lower or very high internal densities (i.e. k-density) of the rich-club. As the rich-club is not dominated by a single or a few functional networks, we subsequently investigated whether instead there are networks that ‘dedicate’ relatively more nodes to the rich-club than other networks. We addressed this by quantifying how many brain regions of a functional network participate in the global rich-club adjusted by the size of the rich-club. Specifically, we calculated the relative network dedication (see Methods). Despite increasing k-densities, the functional networks always dedicated roughly the same fraction of nodes to the global rich-club (Fig. 4b). Whereas, the somatomotor network dedicated the largest and the auditory network the smallest fraction, all other networks showed roughly a similar relative network dedication. These observations characterise this rich-club as a global structural foundation for possible functional network integration.

**Fig. 4:**
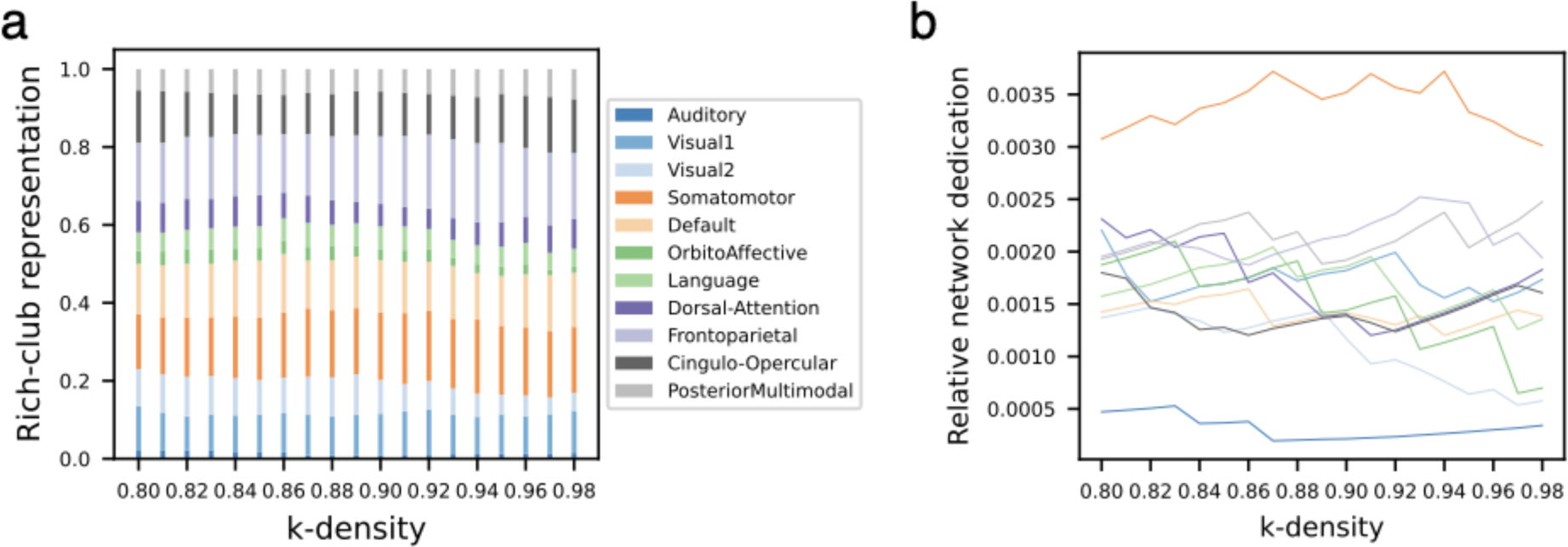
Functional network representation and dedication in the global rich-club. **a** Rich-club representation for different k-densities of the global rich-club. The rich-club representation indicates to which degree a network is represented in the rich-club. Nearly all functional networks participated in the global rich-club (except the ventral multi-modal network) and no clear dominance of a single network can be observed. **b** Relative network dedication for different k-densities of the global rich-club. The relative network dedication indicates to which degree a single network “dedicates” its own nodes towards the rich-club. For increasing k-densities, the functional networks dedicated approximately the same fraction of nodes to the global rich-club. Whereas the somatomotor network relatively dedicated the most of its nodes, the auditory network dedicated the least.

### Signal propagation along the global and local structural (sub)networks crucially depend on hub nodes

Structural network disruptions are hallmarks of various pathologies such as disorders of consciousness (e.g. coma), traumatic brain injury, stroke, and Alzheimer’s disease. The question arises, when are brain lesions especially disruptive? From a structural perspective, the targeted removal of hub nodes has been found to be detrimental to the structural integration of the cortex (Kaiser et al., 2007).

Here we have shown that subcortical rather than cortical hubs display a predominant reach in the global structural network. Whereas hubs were found not only in the cortex but in all areas, subcortical hubs had an especially wide reach in the global structural network. To elucidate how hub lesions disrupt the brain-wide communication, we investigated how targeted lesions affect signal propagation along the global and individual (sub)networks. For that, we employed a model-based network analysis approach which captures the temporal evolution of the network responses to unit perturbations (Gilson et al., 2019). These responses encompass the influence of one node over another throughout paths of different lengths, at different times. In particular, we measured the global network response, r, accumulated over time (from the perturbation onset at t = 0) until the elicited network activity vanishes (*t* → ∞).

First, considering only the global network, we lesioned either all brain regions in a single sequence or only cortical, subcortical, left, and right cerebellar brain regions (Fig. 5a). Importantly, for all sequences we started with the most connected brain region (i.e. most influential hub) and subsequently lesioned the next most connected brain region, etc.. For initial hub lesions (e.g. 50 nodes; red vertical line), exclusive subcortical lesions were most harmful leading to a higher loss in global network response than exclusive cortical or (left and right) cerebellar node lesions (Fig. 5a). The exclusive subcortical hub lesions are also to a large extent as harmful as considering all hubs in a single lesion sequence (global targeted lesions: first 50 nodes also entailed nine cortical hubs).

**Fig. 5:**
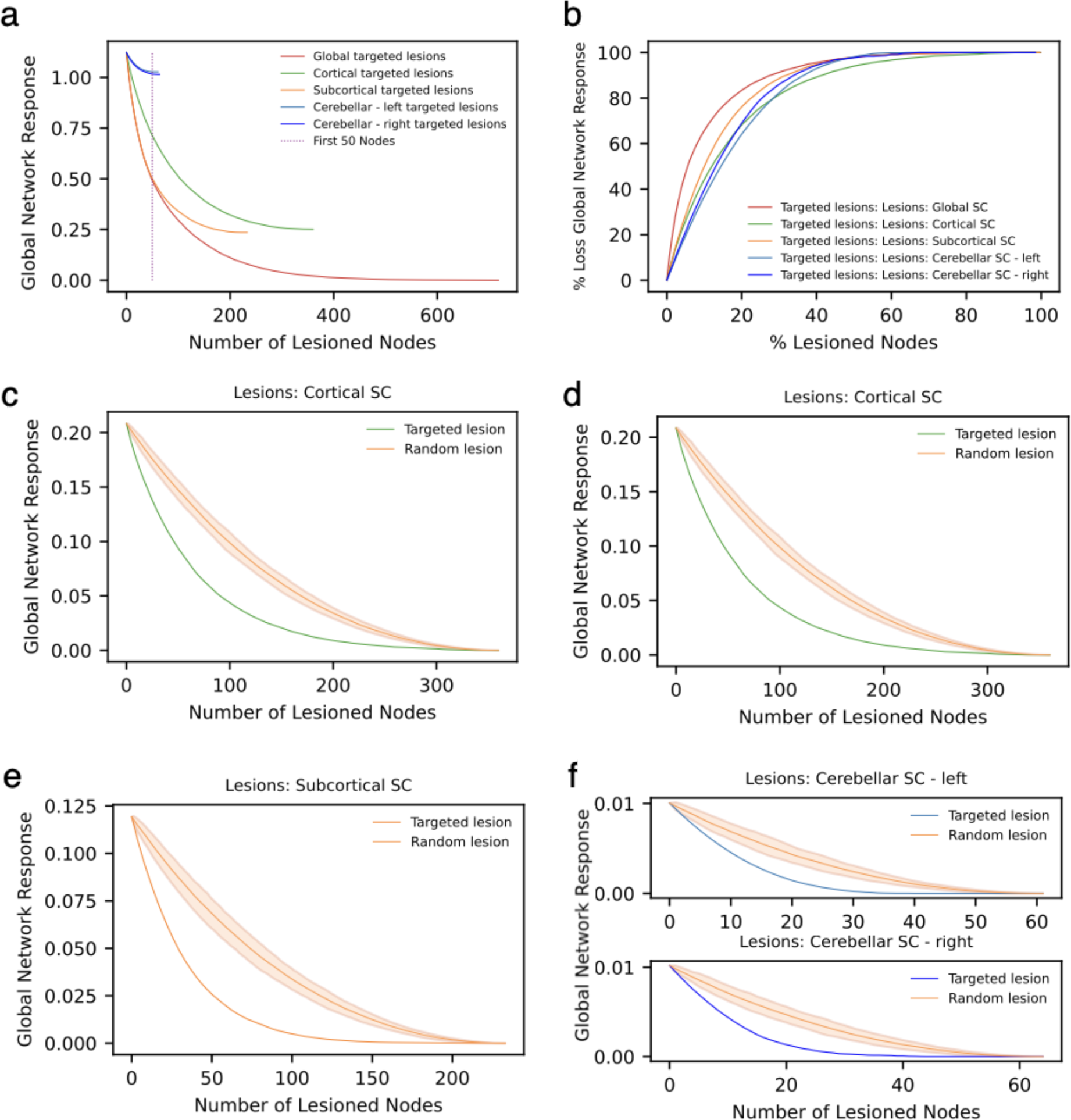
Loss in global network response after targeted and random lesions. **a** Targeted lesion sequence in the global network only. Either cortical, subcortical, and (left & right) cerebellar nodes or all together (Global targeted lesions) were sequentially lesioned. The targeted lesions started with the most connected node and sequentially lesioned the next most connected node. The red vertical bar indicates the first 50 lesioned nodes. Considering brain regions individually, subcortical hub lesions were most harmful. **c-f** Targeted and random lesioning for all individual (sub)networks. The random lesion sequence consisted of 100 different iterations. The mean is illustrated as the solid orange line and the shaded area represents two standard deviations. **b** To compare the loss in global network response between the different (sub)networks, the percentage loss in initial global network response compared to the percentage of lesioned nodes is represented. Initial hub lesions were most harmful to the global network.

Next, we also investigated the sequenced targeted hub removal in the individual (sub)networks and compared these with 100 random lesion sequences. All (sub)networks showed a significant decrease in global network response when the (sub)networks were lesioned targeted compared to random lesions (Fig. 5c-f). These findings generally confirm the importance of hubs for signal propagation in the (sub)networks. For the sake of completeness, we repeated the targeted hub and random node removal considering the classical network’s efficiency (Supplementary Materials S3 a-e). Equivalently, targeted hub lesions were most harmful to the (sub)networks’ capacity to efficiently transmit information compared to random lesions. To enable a comparison between different structural (sub)networks, we compared the relative loss in global network response after targeted lesions (Fig 5b). Interestingly, initial hub lesions were most harmful for the global network highlighting that these lesions were especially detrimental when considering the connectome as a whole, whereas the individual cortical, subcortical, and (left and right) cerebellar subnetworks were more resilient. The discrepancy in global signal loss and for the individual (sub)networks suggests that links between the cortex, subcortex, and cerebellum substantially contribute to global signal propagation.

### Common and distinctive network properties of the cortical, subcortical, and cerebellar subnetworks

Next to a system-level analysis of how the global connectome is structurally integrated, we lastly examined the individual network characteristics of the global network and the cortical, subcortical, left, and right cerebellar subnetworks. We observed both similarities and distinctive architectural principles among them.

The global network was characterised by a broad degree distribution showing a higher prevalence of sparsely compared to densely connected brain regions (Fig. 6a). Similarly, albeit less pronounced, the individual cortical and subcortical subnetworks also showed a broad degree distribution. Notably, the left and right cerebellar subnetworks were characterised by a more uniform distribution, indicating a similar prevalence of differently connected brain regions (Fig. 6b).

**Fig. 6:**
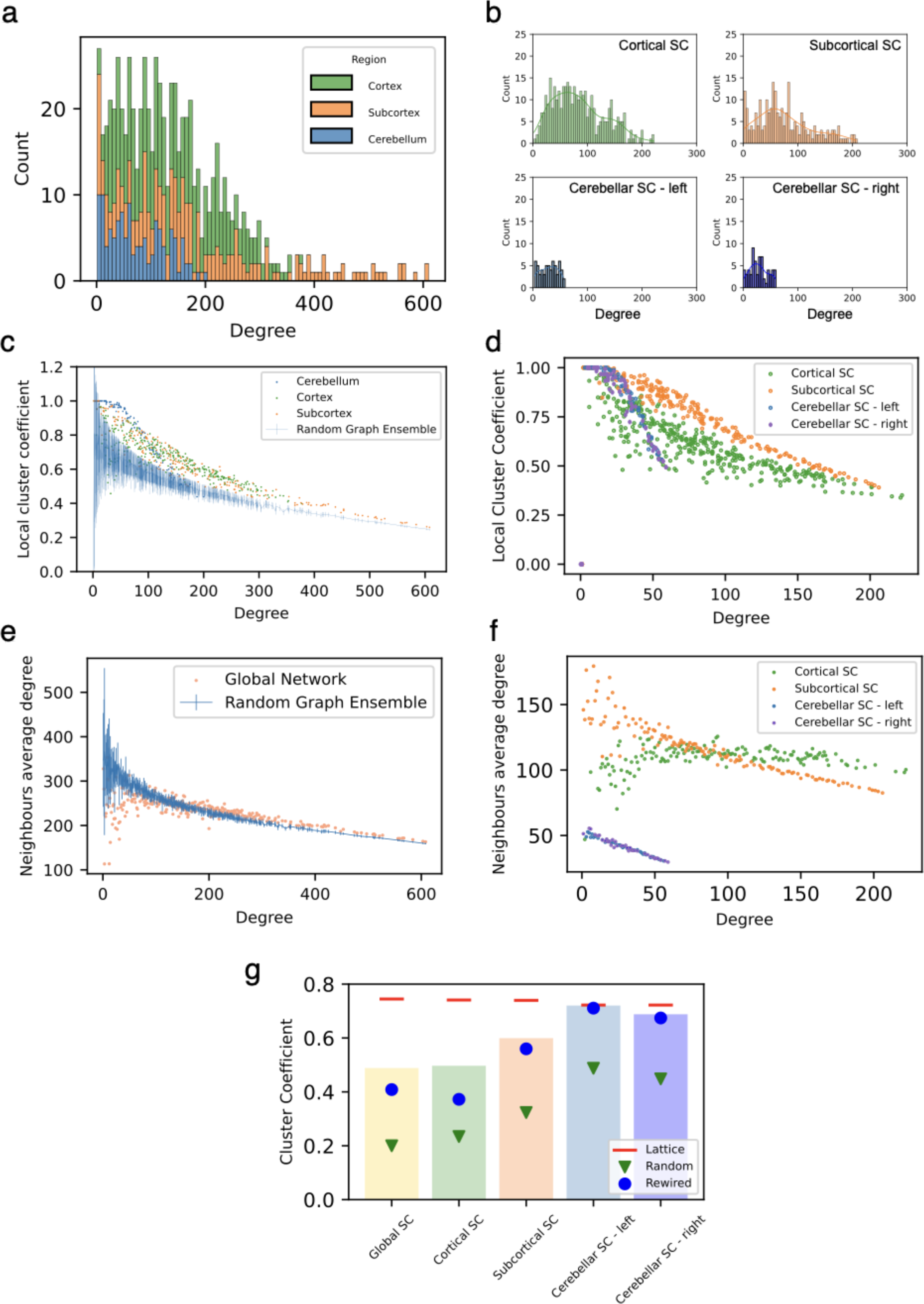
Network description of all (sub)networks. **a,b** Degree distributions of the global (a) and of the individual cortical, subcortical, left, and right cerebellar subnetworks (b). Whereas the global, cortical, and subcortical subnetworks showed a more broad degree distribution with a few hubs and more less connected nodes, the left and right cerebellar subnetworks showed a more uniform distribution. **c** Local cluster coefficient of all ROIs in the global network. The blue line indicates the mean of the random graph ensemble (n=1000) and the error bars two standard deviations. The majority of ROIs were more clustered than expected from the network’s degree distribution. **d** Local cluster coefficients of the individual cortical, subcortical, left, and right cerebellar subnetworks. **e** Neighbor’s average degree of the global network. Generally, a disassortative trend can be observed. **f** Neighbor’s average degree of the individual cortical, subcortical, left, and right cerebellar subnetworks. Whereas for the subcortical, left, and right cerebellar subnetworks disassortative characteristics can be observed, the cortical subnetwork neither exhibited assortative nor disassortative characteristics. **g** The global cluster coefficient of all (sub)networks compared to a lattice, random, and rewired graph with the same size and density. All (sub)networks had a higher cluster coefficient than their random equivalent and nearly all were higher than their rewired equivalent.

To assess how well brain regions are locally clustered, we compared the local cluster coefficient *C*(*i*) of all brain regions with the mean local cluster coefficient of a random graph ensemble. The random graph ensemble consisted of 1000 randomly rewired networks, thus maintaining only the original degree distribution of the network but otherwise losing any other previous architectural feature. In the global network, it can be observed that most cortical, subcortical, and cerebellar brain regions had a higher clustering than what was expected from the ensemble (Fig. 6c). Especially cerebellar ROIs were highly locally clustered with some regions’ neighbours being completely connected to each other. Looking at the individual cortical, subcortical, and cerebellar subnetworks, the local cluster coefficient of individual nodes also generally decreased with increasing density (Fig. 6d). Here, subcortical nodes were generally locally more clustered than cortical nodes. Analysing also the global cluster coefficient, the global, cortical, subcortical, left, and right cerebellar (sub)networks were characterised by generally high and increasing coefficients, surpassing the benchmark coefficients of the random network and partly of the re-wired network (Fig. 6g). Notably, the (sub)networks differed regarding their proximity to the various null-models thus indicating different internal architectures.

A high level of structural integration was observed by examining the (sub)networks’ low average shortest path lengths *l*(*A*)and high efficiencies *E*(*A*). All (sub)networks showed a near minimum shortest average path length given their sizes and densities (Fig. 7a). This structural feature of close nodal proximity is generally referred to as small-world property. Congruently, all (sub)networks had also a near-maximal possible network efficiency (Fig. 7b), which underlines the structural capability of all networks to efficiently exchange information.

**Fig. 7:**
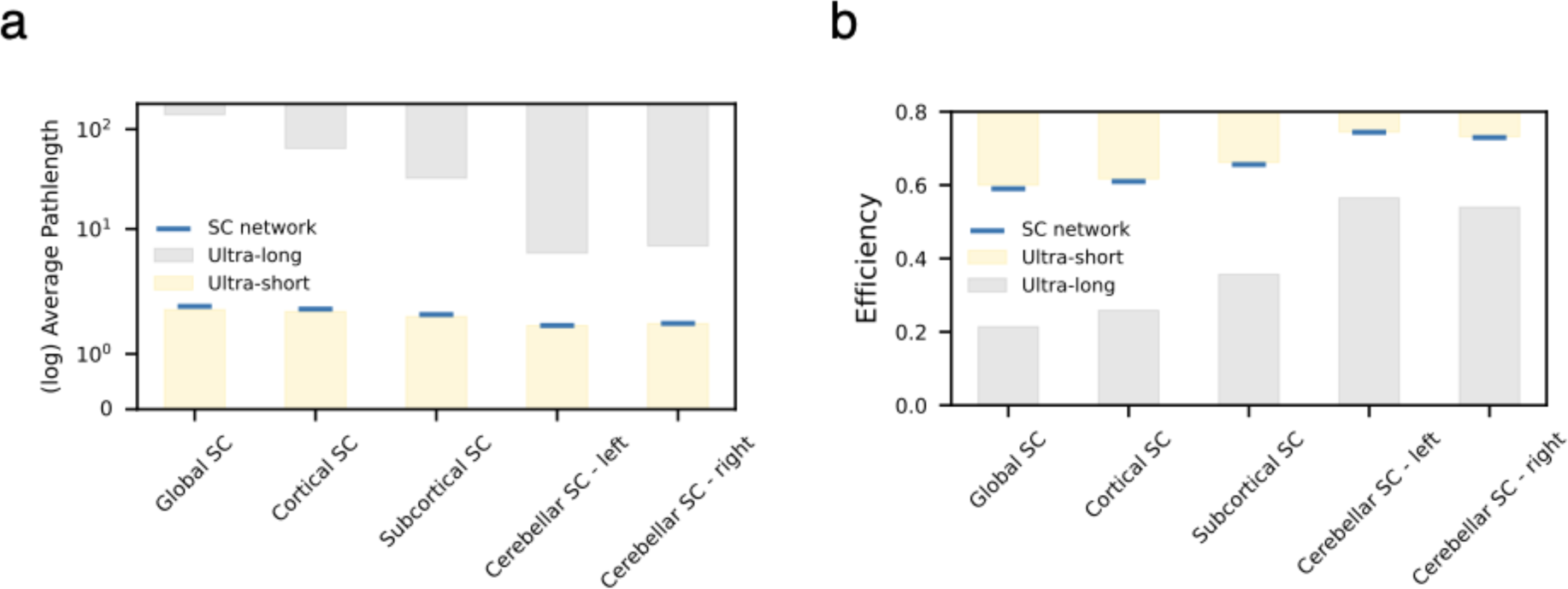
**a** The average shortest path length of the global, cortical, subcortical, left, and right cerebellar (sub) networks (blue). Given the networks’ sizes and densities, the theoretical upper (grey) and lower (gold) limits are displayed. All (sub)networks are close to their theoretical optimum. **b** The efficiency of the global, cortical, subcortical, left, and right cerebellar (sub)networks (blue). Here the theoretical upper (gold) and lower (grey) limits are also represented. All (sub)networks are close to the theoretical optimum.

Structural heterogeneities between the cortex, subcortex, and cerebellum were pronounced when considering the average nearest neighbour’s degree ⟨*k_nn_*(*k*)⟩ (Fig. 6e,f). Higher degree nodes in the subcortical subnetwork connect on average to smaller degree nodes and vice versa (assortativity coefficient *r* = −0.30; Fig. 6f), thus exhibiting a so-called disassortative network characteristic. This property is even more pronounced for the left and (assortativity coefficient *r* = −0.39) right cerebellum (assortativity coefficient *r* = −0.41) and can also be observed for the global structural network (assortativity coefficient *r* = −0.18; Fig. 6e). Notably, the cortex by itself neither appears to be an assortative nor disassortative network (assortativity coefficient *r* = −0.02). Whereas for the left and right cerebellar subnetworks the level of assortativity can largely be explained by their degree distribution (supplementary materials S2), the global, cortical, and subcortical (sub)networks showed for larger degree nodes a higher assortativity than expected (Fig. 6e & Supplementary Materials S2). Consequently, despite similarities in the (sub)networks’ near optimal short-average path length and capacity to transmit information, differences regarding their degree distribution, clustering, and assortativity suggest distinctive architectural features.

## DISCUSSION

The brain’s structural connectivity constitutes the skeleton for inter-areal communication between brain regions and structural damage to this complex network can have detrimental effects on brain health and functioning. Despite this importance, it has remained an open question how collectively cortical, subcortical, and cerebellar brain regions shape the overall network architecture. Here, we found highly dense communities (i.e. communities 1-4) that span over the brain-wide structural network. These transversely cross over cortical, subcortical, and cerebellar boundaries. All five communities are structurally centralised by a global rich-club, which at its core (i.e. high densities) is subcortically dominated, where the remaining fraction is formed by cortical hubs. In particular, subcortical hubs revealed themselves as the most far-reaching brain regions, which at the same time uniformly distribute their links between all the communities and therefore crucially support structural integration. This result is mirrored in the virtual lesioning study, where subcortical hub removal is most detrimental to global signal propagation. These findings highlight the need to shift from a cortex-centric perspective to a brain-wide consideration of the connectome and all its subnetworks.

Considering the entire connectome not only allows for a global analysis of the brain’s network properties but we also investigated the individual cortical, subcortical, left, and right cerebellar subnetworks. Whereas major attention has been historically devoted to the network analysis of the cortex, and rarely included subcortical regions, a side-by-side comparison also accounting for the cerebellar subnetwork has been missing until this point. We find that despite large commonalities shared between these subnetworks, unique structural fingerprints were detected. All (sub)networks, including the global structural network, exhibit a near-optimal average path length and efficiency. Distinctive differences could be observed in the degree distributions, clustering, and assortativity. These unique fingerprints highlight that also subcortical and cerebellar network properties should be regarded when aiming to understand the brain from a system-level perspective.

From a neurobiological point of view, it is well known that subcortical regions for instance the amygdala (Phelps & LeDoux, 2005), the hippocampus (Huang et al., 2021; Maller et al., 2019), the basal ganglia (Alexander et al., 1986; Haber & Calzavara, 2009), and the thalamus (Castro-alamancos & Connors, 1997; Haber & Calzavara, 2009; Jones, 2001) are heavily interlinked with the cortex via a multitude of pathways. Additionally, the cerebellum is interlinked to the cortex via the cortico-ponto-cerebellar and the cerebello-thalamo-cortial pathways (D’Angelo, 2018; Palesi et al., 2015, 2017; Ramnani, 2006). This tight interconnectedness between cortical, subcortical, and cerebellar brain regions is emphasised by the detected communities. Whereas modular properties have been previously shown for the cortex alone (Betzel et al., 2013; Hilgetag et al., 2000; Hilgetag & Kaiser, 2004; Newman, 2004; Zamora-López et al., 2011), communities one to four transversely stretch across cortical, subcortical, and cerebellar boundaries and show equally or even higher internal connection probabilities than in the cortex, subcortex, and cerebellum. Interestingly, brain regions in communities three and four show a hemispheric lateralization thus potentially supporting as well more lateralized functions such as language (Knecht et al., 2000; Olulade et al., 2020) and attention (Bartolomeo & Seidel Malkinson, 2019). The other communities are represented in both hemispheres and show a more medial distribution thus possibly supporting more integrative processes with regards to both hemispheres.

Overall structural integration is achieved by a global rich-club integrating the pathways between the communities. For the cortex alone, this centralising rich-club property has been already established (van den Heuvel & Sporns, 2011; Senden et al., 2014, 2017, 2018; Zamora-López, 2010; Zamora-López et al., 2011) and we find this architectural property is also represented in the global connectome. At lower rich-club densities, the occurrence of subcortical and cortical hubs is roughly equal, however when considering the core of the rich-club (i.e. at highest k-densities), a clear dominance of subcortical rich-club nodes can be observed. Additionally, we find that numerous subcortical hubs are the most far-reaching brain regions in the global network with highly integrative structural properties. Taking these findings together strongly emphasise the centralising role of subcortical brain regions. Congruently, our subsequent lesioning study revealed that subcortical hub lesions, compared to cortical or cerebellar hubs are most harmful to signal propagation along the global network. Although these results only reflect a reduced signal propagation caused by structural damage and no consequences for functionality can be directly inferred, damage to subcortical brain regions has been for instance shown to be detrimental to loss of consciousness (Fischer et al., 2015; Panda et al., 2023.).

The global rich-club not only facilitates structural integration but might also serve as an anatomical scaffold for functional integration. Previous research has suggested this notion for the cortex individually, as it has been observed that for instance all functional networks are represented in the cortical rich-club (van den Heuvel & Sporns, 2013). Expanding this analysis to the global network, we confirm that nearly all major functional networks (except the ventral multimodal network) participate in the global rich-club and that all networks approximately dedicate the same fraction of nodes to the rich-club. To this point, it is not clear how a functional integration via the rich-club could be mechanistically obtained. However, previous research found that the existence of a rich-club architecture promotes a network’s heterogeneity of functional repertoire and might achieve functional integration by aligning asynchronous brain regions via oscillations (Senden et al., 2014, 2017).

This study has several limitations. The found architectural principles of the structural network are bounded by the resolution and technical possibilities of the DWI. Therefore, we described the general properties of the structural network regarding the overall cortical, subcortical, and cerebellar regions. Future studies employing a higher imaging resolution could enable more detailed analysis and interpretation even between the individual ROIs. Moreover, a general limitation is the occurrence of false-negative and false-positive connections when estimating the structural connectivity based on DWI. The current DWI data was already pre-processed and previously a generalised q-sampling imaging algorithm (Yeh et al., 2010) together with the Gibbs Tracking approach (Kreher et al., 2008) were applied, which aim to account for the crossing and spreading of fibres. To further reduce the risk of false-negative connections, we thresholded the connectivity matrix such that the initial length distribution was preserved and therefore long-range connections were maintained. However, as some ROIs had only a few short-range connections with low connection weights, we assured that their strongest connections survived, leading to a minor adjustment of 12 additional links. As a last effort to reduce the presence of non-existing direct structural connections, direct cortico-cerebellar links, interhemispheric links in the cerebellum, and ipsilateral cerebellar connections to the thalamus and brainstem were filtered out, as these do not reflect previous anatomical knowledge (D’Angelo, 2018; Palesi et al., 2015, 2017; Ramnani, 2006). Despite these precautions, the extent of false-positive and false-negative connections is not known due to the nature of DWI. Therefore, this study represents the first step toward a global understanding of the structural network, and further studies have to come applying more refined methods. Despite the current limitations of the dataset, it is still important to investigate the global structural connectome and to find common and distinctive network properties of the (sub)networks to foster an overall understanding of the connectome. The upcoming availability of high-resolution connectomes in the future will aid this quest. We are confident that the general qualitative results reported here will remain despite potential quantitative differences.

Additionally, different brain parcellations could also be applied to investigate the structural connectome from further perspectives. In detail, more unified parcellations that encompass cortical, subcortical, and cerebellar areas altogether could aid future investigations. Lastly, as many whole-brain computational models are constrained by the structural connectivity and mainly focus on cortical brain areas, the inclusion of subcortical areas, especially considering their centralising importance, might further facilitate the understanding of whole-brain dynamics given appropriate models. Despite these limitations and future studies to come, we established with these results a first important step towards a whole-brain understanding of the global architecture and how structural integration is facilitated by its unique characteristics and fingerprints.

## METHODS

### Subjects and dataset

The current dataset is from Ji et al. (2019) and taken from the Human Connectome Project (HCP) with 32 participants collected at the Massachusetts General Hospital (“MGH HCP Adult Diffusion,” 16 females, 16 males; Van Essen et al., 2013).

### Data Acquisition

The whole-brain echo-planar imaging was conducted with a modified 3T Siemens Skyra (Connectome Skyra) system with a 32 channel head coil applying a time to repetition (TR)=720ms, time to echo (TE)=33.1ms, flip angle=52, bandwidth=2,290 Hz/pixel, in-plane field of view (FOV)=208×180mm, 72 slices, and 2.0mm isotropic voxels, with a multi-band acceleration factor of 8 (Uğurbil et al., 2013). For more details see Ji et al. (2019).

### Structural scan

The structural connectomes of the participants were averaged to generate a representative connectome. For the DWI a high-quality protocol was applied (i.e. b-value of 10,000 s/mm2, high-angular resolution, and a multi-slice approach). The data are publicly included within the Lead-DBS software package and were already preprocessed (Horn et al., 2017; Setsompop et al., 2013). A generalised q-sampling imaging algorithm (DSI Studio; http://dsi-studio.labsolver.org) was used for the data processing. SPM 12 was used to segment and co-register the data. 200 000 fibres were examined for each participant using Gibbs’ tracking approach (Kreher et al., 2008) and being restricted by a co-registered white-matter mask. The fibres were standardised into MNI space via DARTEL transforms (Ashburner, 2007; Horn & Blankenburg, 2016). The cortex was parcelled into 360 regions (180 per hemisphere) according to Glasser et al. (2016).

### Functional networks & subcortical/ cerebellar ROI assignment

The functional network calculation and the subcortical/ cerebellar ROI assignment was previously performed by Ji et al. (2019). In detail, subject-specific functional connectivity matrices were calculated by correlating the individual cortical BOLD signals with Pearson correlation. By averaging across all subject matrices, a group-averaged functional connectivity (FC) matrix was constructed. Cortical resting-state functional networks were obtained by applying the Louvain algorithm to this group-averaged functional connectivity matrix (Blondel et al., 2008). Next, subject-wise FC matrices indicating the correlation between the 360 cortical ROIs and 31870 subcortical/ cerebellar grayordinates spanning over the entire CIFTI space were calculated and averaged to obtain a group-representative cortical - subcortical/ cerebellar FC matrix. These grayordinates were assigned to cortical functional networks with the highest correlation and merged together based on this partition. At the same time, the parcels were constrained to major subcortical structures as determined by Freesurfer, thus adhering to the general subcortical anatomy. This resulted in a partition of 233 subcortical and 125 cerebellar ROIs based on the detected cortical functional networks (for details see Ji et al., 2019).

### Filtering false-positive structural connections

The cortex and cerebellum are contra-laterally connected via the cortico-ponto-cerebellar (CPC) and the cerebello-thalamo-cortical (CTC) pathways (D’Angelo, 2018; Palesi et al., 2015, 2017; Ramnani, 2006). To only display direct anatomical connections, initial cortical-cerebellar connections were filtered as diffusion imaging cannot distinguish between direct and indirect pathways. This resulted in the filtering of 8.3% of initial connections. Moreover, ipsilateral connections between the cerebellum and the brainstem (pons)/ thalamus (total of 2.0% of initial connections) and direct inter-hemispheric cerebellar connections (as the cerebellar hemispheres are indirectly connected via the vermis (D’Angelo, 2018); 2.6% of initial connections) were filtered.

### Binarization of the population-level connectivity matrix

The individual SC matrices for the 32 participants were averaged into one population brain-wide structural connectivity network, represented as the weighted global connectivity matrix *W* of *718 × 718* (ROIs). To perform a graph analysis, commonly *W* is binarised by applying a hard threshold *θ* such that only the connections with a weight larger than *θ* are conserved. That is, the binary adjacency matrix *A* is defined as *A_ij_*= 1 if *W_ij_* > *θ*, and *A_ij_* = 0 if *W_ij_* < *θ.* The value of *θ* then controls the final number (density) of connections. This typical hard thresholding leads to a bias. Since one of the limitations of tractography is that it overestimates shorter connections, the hard thresholding favours short-range links in detriment of long-range ones (Betzel et al., 2019; Roberts et al., 2017). Nevertheless, the existence of long-range connections has been shown in tract-tracing studies (Ercsey-Ravasz et al., 2013; Horvát et al., 2016) and are thought to serve important functional roles (Betzel et al., 2019) as these connections cause a higher metabolic cost for the organism (Bullmore & Sporns, 2012). To threshold the weighted matrix *W* while also maintaining long-range connections, we introduce the “*relative weight distance-dependent*” (RWDD) thresholding. The goal of RWDD is to conserve the original distribution of connection lengths by performing an adaptive threshold such that *θ = θ_d_* depends on the distance *d(i,j)* between two ROIs. Here, *d(i,j)* will represent the Euclidean distance between the centers of mass of two ROIs *i* and *j*, but it could be replaced by the fiber-length of the tracts when this information is available.

The weighted connectivity W is an all-to-all connected matrix. All links of the initial SC matrix are binned into *N* bins according to their length. This information is taken from a distance matrix, representing the distances between all nodes. Thus, each bin contains similarly long connections. Next, the relative weight of each bin is calculated i.e. the number of connections in the bin compared to all connections in the initial SC matrix. Finally, the bins are reduced such that this relative weight is maintained and the number of final links for the thresholded SC matrix is reached. While doing so, only the strongest connections of the bins are retained.

Consequently, the RWDD thresholding was applied with a target density of 0.2. While targeting for a low density to reduce false-positive connections, unconnected components (i.e. ROIs with no connections to other ROIs) could be created. To account for this, strong connections of otherwise unconnected ROIs were being kept. If an ROI had the risk of becoming disconnected, links with a weight of 1.8 standard deviations above the mean were kept. This resulted in 12 additional links conserved which would initially violate the RWDD thresholding criteria.

### Network analyses

Several network measures were applied to characterise the global, cortical, subcortical, left and right cerebellar (sub)networks.

### The density

*ρ*(*G*) of a network, which is the fraction of its existing number of links and the maximum possible number of links 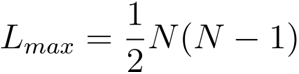, where is the number of

ROIs in a network. Similarly, the connection densities between areas in a network (e.g. cortex and subcortex) were calculated as the fraction of existing links *L_region_* between these areas and the maximum possible number of links *L_maxregion_* = *N_i_ N_j_* with *N_i,j_* the size of areas *i* and *j*.

### Global hubness

The hubness *h_i_* compares the degree *k_i_* of a node with the degree distribution of a corresponding random network with an equal size *N* and density *ρ* (Klimm et al., 2014), thus creating a common random benchmark:

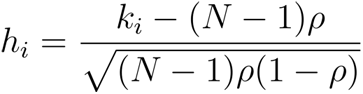

### The participation coefficient

*p_i_* is a measure of centrality and assesses the relative importance of nodes (Rubinov & Sporns, 2010). Here we applied the participation coefficient proposed by Klimm et al. (2014) as it takes into account differently sized communities. A participation coefficient of zero indicates that node *i* has only links within the own community, whereas a value of one describes a uniform distribution of links towards all communities. It is defined as:

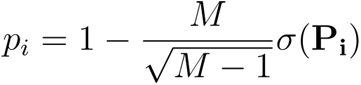

where M is the number of communities, **P_i_** the participation vector of node *i* indicating the probability that *i* belongs to community *C_m_*, where *m* = 1,2…, *M* and σ(**P_i_**) is the standard deviation of **P_i_**.

### A Rich-club

consists of highly interconnected hubs (Zhou & Mondragon, 2004). It is iteratively calculated by only considering nodes with a degree larger than *k′* resulting in networks with a k-density *ϕ*(*k′*):

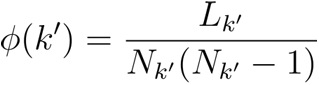

Where *N_k′_* is the number of nodes with *k*(*v*) > *k′* and *L_k′_* the number of links between these nodes (Zhou & Mondragon, 2004). As rich-club nodes can be identified for different k-densities, a range of *ϕ*(*k′*)= [0.8, 0.98] was considered. We compared the k-density of the global, cortical, subcortical, and cerebellar (sub)networks with an ensemble of 1000 randomly re-wired networks with the same densities. To evaluate the extent to which resting-state functional networks participated in the global rich-club, we measured the rich-club representation (RCR) as the fraction of network specific ROIs in the rich-club compared to the total number of nodes in the rich-club. To also understand how much a network ‘dedicates’ of its nodes to the global rich-club while adjusting for the size of the rich-club, we defined the relative network dedication (RND) as:

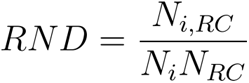

Where *N_i,RC_* are the number of nodes in network *i* participating in the rich-club, *N_i_* is the total size of network *i* and *N_RC_* is the size of the rich-club.

### The global clustering coefficient

*C* measures the conditional probability that any two nodes are connected given they have a common neighbour. It is calculated by comparing the number of triangles in the network *N*(∇) with the number of paths with the length of 2 *N*(V) (Wasserman & Faust, 1994):

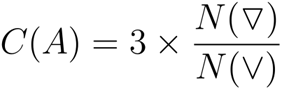

### The local cluster coefficient

*C*(*i*) provides information about how densely node i’s neighbours are connected in adjacency matrix *A* for the (*i, j*)th element *a_i,j_* (Wang et al., 2017):

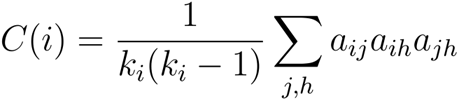

where *k_i_* is the degree of node *i*.

### The average nearest neighbour’s degree

⟨*k_nn_* (*k*)⟩ by Pastor-Satorras et al. (2001) was evaluated by averaging a node *i*’s neighbours degrees, *k_nn, i_*, and then averaging over all nodes with the same degree as *i.* To assess the degree of assortativity, we applied the assortativity coefficient *r* (Newman, 2002). A positive coefficient indicates an assortative network, whereas a negative coefficient indicates a disassortative network.

The **average shortest path length** *l* was calculated as:

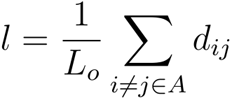

where *d_ij_* is the shortest distance between and and N the number of nodes in A and *L_o_* = *N*(*N* − 1) is the maximum possible number of links.

### The network’s efficiency

*E* (*A*)(Latora & Marchiori, 2001) was also assessed:

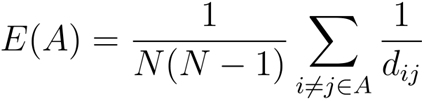

The lower and upper limits for the average path lengths and efficiencies for the individual networks were subsequently calculated (Zamora-López & Brasselet, 2019).

### Community detection

Generally, a community is concerned as a subset of nodes, which are more densely interconnected inside the community than to nodes outside of that community (Girvan & Newman, 2002). Here, we applied the Leiden community detection algorithm (Traag et al., 2019) for the global SC matrix. To account for different resolution parameters, we applied in a first step the RB null-model proposed (Reichardt & Bornholdt, 2006). For each step of size of 0.05 in the range of γ ∊ [0.6..1.9], 100 partitions were calculated and the partition with the highest quality value was selected. Finally, the partition of this set with the highest modularity index (Newman & Girvan, 2004) was chosen.

### Randomly rewired graph ensemble

To destroy the internal architecture of the graph while only preserving the degree distribution, we applied a link-switching model (Kannan et al., 1999; Rao et al., 1996; Snijders, 1991). Zamora-López (2009) showed that 1 x L iterations are sufficient to destroy the internal network architecture, where L is the number of links. Here one random network was iterated 10 x L times to ensure randomization. In total, 1000 random networks for the individual global, cortical, subcortical, and cerebellar (sub)networks were in this way constructed and used for further significance testing.

### Random and lattice benchmark networks

We compared the global clustering of the (sub)networks also with a random and lattice benchmark networks with the same sizes and densities. The random graphs were calculated according to the random generator known as the G(*n,m*) model which ensures the same number of links. For the ring lattice network, each ROI is linked to its closest neighbours, therefore ensuring a high clustering and that all nodes have the same degree.

### Targeted and random network lesions

Nodes were removed from the adjacency matrix A such that the row and column entries of the nodes were set to zero. For the global network response propagation, the entire rows and columns were removed from the adjacency matrix to increase computational efficiency.

### Perturbation-response network analysis

We employed a model-based approach to quantify the effect of random and targeted lesions on the networks. This approach captures the influence between two nodes through all paths of all lengths (Gilson et al., 2019), beyond the traditional procedures that account only for the effects through shortest paths. Considering the continuous propagation model of activity

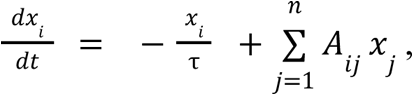

the term 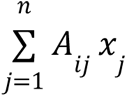 represents the actual propagation of activity across the connectivity and 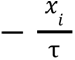 is a leakage or dissipation term representing an exponential decay of the activity at each brain region, with the decay rate determined by τ. For small enough values of τ –say, fast enough leakage – the divergent behaviour of the second term is counterbalanced through this dissipation term. This allows to characterise the transient response of a network to a unit perturbation applied at time t = 0 at all nodes. Given that 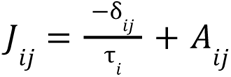 is the Jacobian matrix of the linear system with δ*_ij_* is the Dirac-delta, the pair-wise response function can be defined as:

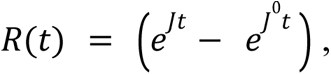

where we regress out the term 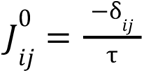 corresponding to the trivial leakage through a region due to the perturbation applied on itself. On the one hand, the pairwise elements *R*_*ij*_ (*t*) represent the conditional, temporal response of area *j* to a unit perturbation applied on area *i* at time *t = 0*. This pair-wise response encompasses all network effects from *i* to *j* acting at different time scales along all recurrent paths of different lengths. On the other hand, at every time point *t*, *R*(*t*) is an *n* × *n* matrix revealing patterns of spatio-temporal interaction that emerge from connectivity A, at time t > 0. Depending on the field of research, the unit response function *R(t)* is often named as the Green’s function (Bayin, 2013) or the propagation kernel.

The evolution of network response *r*(*t*) represents the total amount of pair-wise responses propagated through the connectivity over time. It is computed as the sum of the pair-wise responses *R_ij_* (*t*) at every time point t, 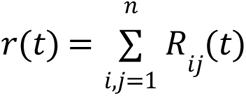. Finally, for every connectivity matrix A, either complete or lesioned, we computed the corresponding global network response *r*, computed as the integral, or area-under-the-curve,

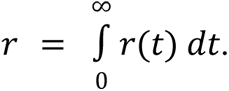

For the results to be comparable across networks, we chose a common and large enough τ for all (sub)networks such that all systems would decay after initial signal growth. Therefore, we set 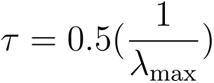, where λ_max_ is the largest eigenvalue of the non-lesioned global connectivity matrix A.

## ACKNOWLEDGEMENTS

We thank Egidio d’Angelo, Claudia Gandini Wheeler-Kingshott, and Fulvia Palesi for the insightful and inspiring recommendations and discussions. This project/research has received funding from the European Union’s Horizon 2020 Framework Programme for Research and Innovation under the Specific Grant Agreement No. 945539 (Human Brain Project SGA3).

## SUPPORTING INFORMATION

Additional supporting information can be found online in the Supporting Information section at the end of this article.

## TECHNICAL TERMS

**Structural connectivity (SC):** Network of white-matter fibres connecting brain regions to each other.

**Hubs:** Highly connected brain regions (here ROIs).

**Community:** Nodes that are are more densely connected to each other than outside of that community.

**Community detection:** Usually an algorithmic process of finding densely interconnected brain regions (i.e. communities).

**Rich-club:** Hubs, which form themselves a highly interconnected community.

**Resting-state network:** Functional community of the brain regions, which occurs consistently between participants at rest (no-task activity).

**In-silico lesion**: Virtually performed lesion to the structural network by either setting the node’s connections to zero or completely deleting the node from the adjacency matrix.

**Perturbation-response network analysis:** A model-based network analysis approach based on the propagation of responses to unit perturbations applied to the nodes.

**Network response:** Temporal evolution of the sum of responses between nodes.

