## Supplementary Materials for "The global communication architecture of the human brain transcends the subcortical - cortical - cerebellar subdivisions"

### SUPPORTING INFORMATION

#### S1: Community participation remains stable over different k-densities

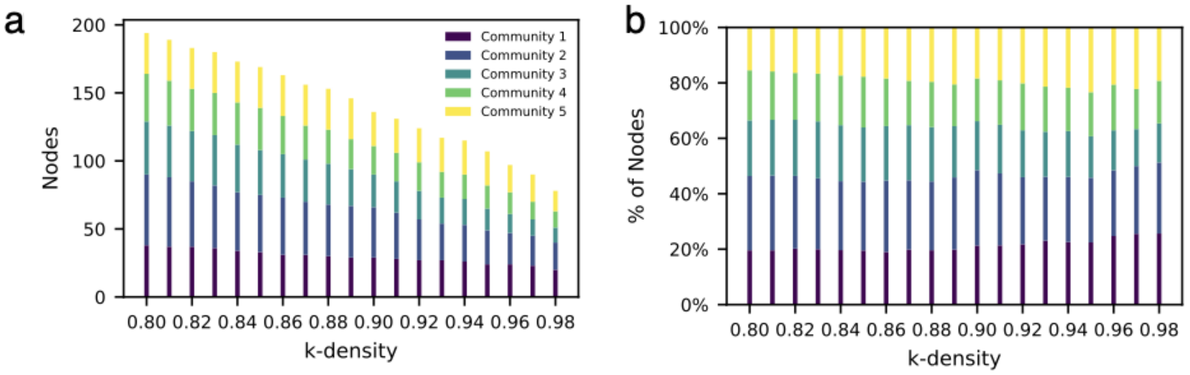


**S1:** Community participation for the rich-club per k-density. **a,b** Total number **(a)** and percentages **(b)** of nodes in the global rich-club per k-density. All communities (1-5) participated in the global rich-club and no clear dominance of a single community can be observed.

#### S2: ANN - Individual subnetworks in comparison with randomly re-wired graph-ensemble

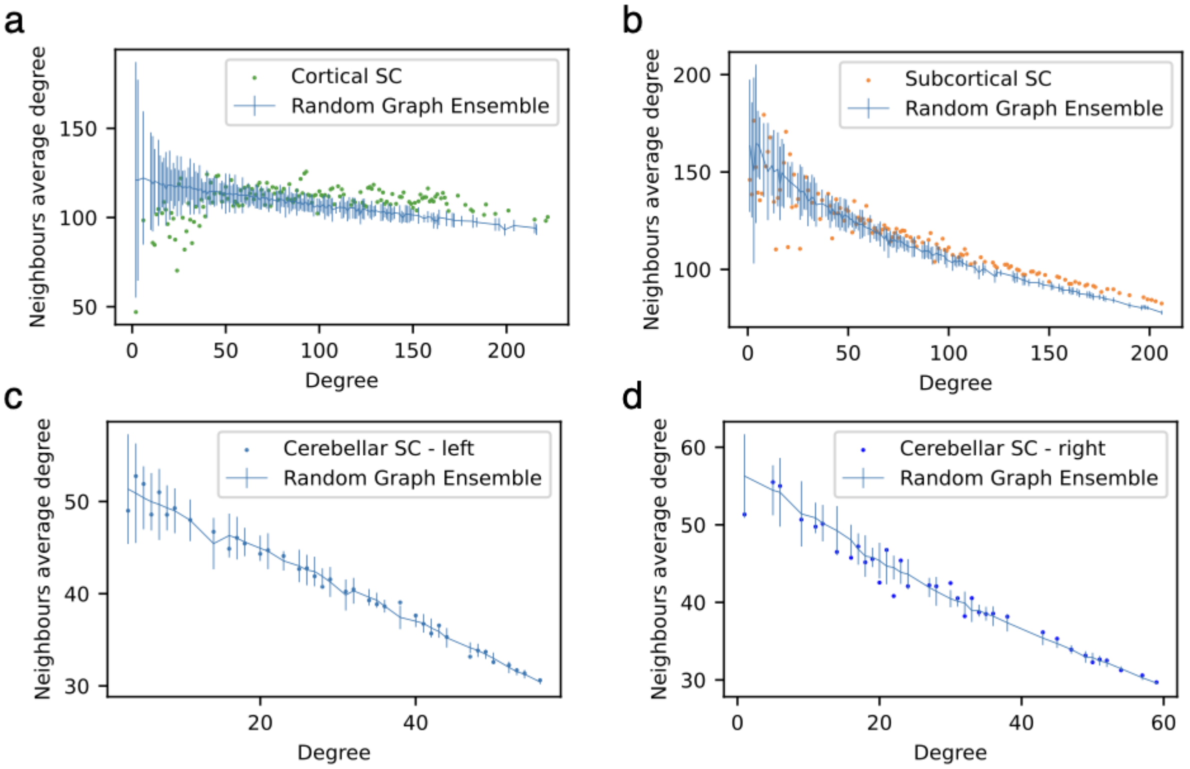

**S2:** Average nearest-neighbour's degree (ANN) for the cortical **(a)**, subcortical **(b)**, left cerebellar **(c)**, and right cerebellar **(d)** subnetworks. The blue line represents the average of a random graph ensemble (n=100) and the error bars are two standard deviations. Whereas, the ANN of cerebellar left and right hemispherical ROIs can be largely explained by the graphs' degree distribution, the cortical and subcortical subnetworks show significant deviations from the random benchmark model.

#### S3: Efficiency: Targeted hubs and random lesion sequences

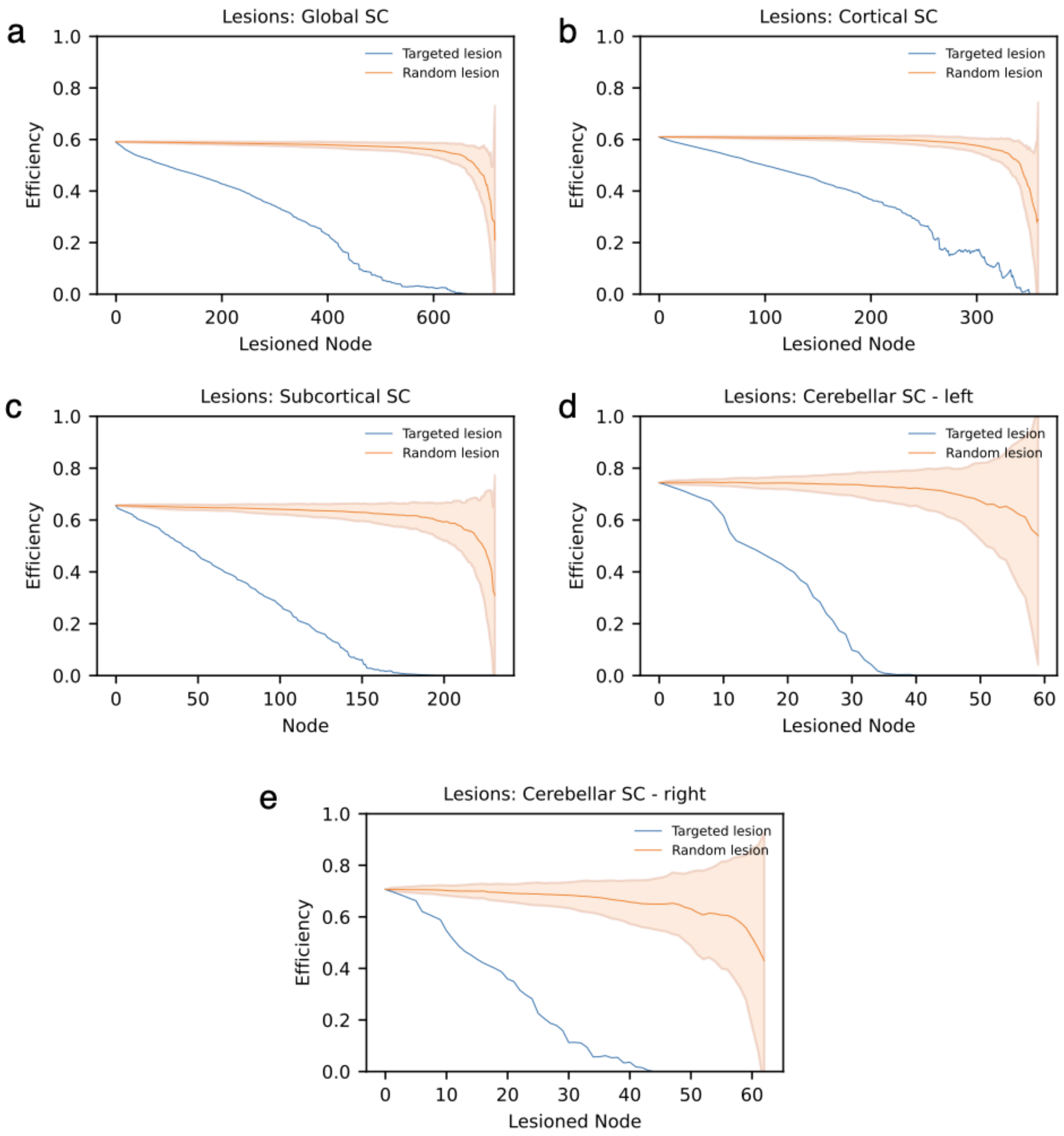


**S3**: Global efficiency of the overall network **(a)**, cortical subnetwork **(b)**, subcortical subnetwork **(c)** , left **(d)**, and right **(e)** cerebellar subnetwork after sequential lesioning starting with the highest degree nodes **(blue line)**. This is contrasted with 100 random lesion sequences **(orange line)**. The orange shaded area indicates two standard deviations above and below the mean. The targeted hub removal resulted in all (sub)networks in a significant decrease in efficiency.
